# No evidence that LSD microdosing affects recall or the balance between distracter resistance and updating

**DOI:** 10.1101/2021.12.02.470935

**Authors:** Sean James Fallon

## Abstract

The effect of low doses (<=20 μg) of LSD on working memory, in the absence of altered states of consciousness, remain largely unexplored. Given its possible effects on serotonin 5-HT_2A_ receptors and dopaminergic signalling, it could be hypothesised that LSD microdoses modulate working memory recall. Moreover, in line with computational models, LSD microdoses could exert antagonistic effects on distracter resistance and updating. Here, we tested this hypothesis in a randomised double-blind, placebo-controlled study comparing three different LSD microdoses (5 μg, 10μg and 20μg) with placebo. After capsule administration, participants performed a modified delay-match-to-sample (DMTS) dopamine-sensitive task. The standard DMTS task was modified to include novel items in the delay period between encoding and probe. These novel items either had to be ignored or updated into working memory. There was no evidence that LSD microdoses affected the accuracy or efficiency of working memory recall and there was no evidence for differential effects on ignoring or updating. Due to the small sample of participants, these results are preliminary and larger studies are required to establish whether LSD microdoses affect short-term recall.

## Introduction

Despite the resurgence of interest in LSD and other psychedelics, the full cognitive effects of LSD remain largely unarticulated (Rifkin et al., 2020). This situation will likely hamper the therapeutic deployment of these substances and in the case of LSD, there is a pressing need to evaluate its cognitive effects in the absence of drug-induced changes in conscious experience. Moreover, the widely reported practice of microdosing – taking very low doses of LSD – to improve cognitive and emotional functioning necessitates an evaluation of the cognitive effect of this practice (Bershad et al., 2019). However, relatively few, laboratory-based, placebo-controlled studies that have evaluated the cognitive effects of microdosing (Hutten et al., 2020; Pokorny et al., 2020). Here, we investigate the effect of very low doses of LSD on working memory.

Working memory is a form of temporally sustained attention to features no longer in the environment that enables us to carry out goal-directed behaviour. Larger working memory capacity is positively related to various academic, economic and health-related outcomes (Gathercole et al., 2005; Kokosi et al., 2021; Rossi et al., 2016; Stautz et al., 2016). However, in addition to the retention of information, working memory systems need to efficiently maintain information in the presence of distracting input whilst enabling representations to be flexibly updated (Braver & Cohen, 2000; Fallon, Zokaei, & Husain, 2016; Myers et al., 2017). The neurocognitive mechanisms through which these multiple demands are reconciled has received considerable attention in recent years (Armbruster et al., 2012; Chatham & Badre, 2015; Dreisbach & Fröber, 2019; Yan & Wang, 2020).

There is a broad consensus that the stability and flexibility of mental representations are reconciled within fronto-striatal circuits (Baier et al., 2010; Chatham et al., 2014; Lewis et al., 2004; Ye et al., 2021; Yu et al., 2013) and that the levels of dopaminergic (either within frontal or striatal circuits) might determine the balance between these two processes (Cools & D’Esposito, 2011; Durstewitz & Seamans, 2008; Hazy et al., 2007). Specifically, computational and empirical evidence has augmented classical notions that working memory recall is dopamine-dependent and requires optimal levels of dopaminergic stimulation (Goldman-Rakic, 1995), with the idea too much or too little dopaminergic signalling can improve working memory where cognitive flexibility is required (Durstewitz & Seamans, 2008; Hazy et al., 2007). Support for this comes from studies that have shown administering D_2_ agonists can differentially affect the stability and flexibility of working memory recall once baseline dependence is considered (Broadway et al., 2018; Fallon, Kienast, et al., 2019). Despite its presumed role in modulating affective functioning, serotonin signalling, particularly 5-HT_2A_ receptors in the prefrontal cortex, also has a role in working memory, i.e., 5-HT_2A_ agonists can disrupt the sustained neuronal delay-period firing (Williams et al., 2002). Therefore, microdoses of LSD, acting either through 5-HT_2A_ receptors or dopamine d2 receptors (De Gregorio et al., 2016), could not only affect overall working memory performance but differentially modulate ignoring and updating. One previous study found no effect of LSD microdoses on performance on the n-back task (Bershad et al., 2019). However, cognitive stability and flexibility are conflated within the n-back and therefore it could be hard to discern whether LSD microdoses had dissociable or antagonistic effects on the stability and flexibility of mental representations. Indeed, there are some reports that low doses of LSD have been found to impair flexibility and working memory (Hutten et al., 2020; Pokorny et al., 2020). However, hitherto, the stability and flexibility of mental representations has not been directly compared.

Here, we examined this question in older adults using a double-blind, placebo-controlled design in which three levels of microdoes (5μg, 10μg, 20μg) were compared with placebo on the performance of a modified delay-match-to-sample task that has previously been shown to be sensitive to pharmacological compounds (Fallon & Cools, 2014). Namely, we modified the standard DMST format (where a probe item had to be identified as matching or not matching the encoded stimuli) to include novel stimuli into the delay period. These novel stimuli either had to be ignored as irrelevant or updated into working memory as the new target stimuli. Thus, we can examine whether LSD affects overall working memory performance or exerts dissociable effects on ignoring and updating.

## Methods

### Participants

The same 48 participants reported in (Family et al., 2020) also took part in this study. Participants were native English-speaking older adults (21 female, 27 male) between 55 and 75 year of age (M = 6.2.92, Sd = 5.65). Recruitment took place from the Early Phase Clinical Unit, Northwich Park, UK. Inclusion criteria were as follows: 1) did not use LSD in the last five years, 2) post-menopausal (if female), 3) male participants with a female partner agreed to use double barrier method of contraception and not to donate sperm for 3 months after last dose. Exclusion criteria: 1) history of psychiatric, neurological, gastrointestinal, immunological, renal, haematological, lymphatic, respiratory, hepatic, cardiovascular, musculoskeletal, genitourinary, dermatological, connective tissue or sleep diseases/disorders and/or intracranial hypertension, impaired consciousness, lethargy, brain tumor, atopy, hypersensitivity, skin allergies or allergic reactions to drugs,2) blood pressure over 160mmHg (systolic) and 90mmHg (diastolic) average across four screening-day assessments, laboratory test results outside the reference ranges; and/or positive results for human immunodeficiency virusm, hepatitis B surface antigen (HBsAg), hepatitis C virus. 3) current smoker of history of drug abuse within last 12 months. 4) prescribed medication or over-the-counter medication (including vitamins with seven days of first doses (unless medical monitor agreed it not clinically relevant), no serotonin-modifying drug (e.g., 5-HT, St. John’s Wort). 5) received or donated blood (3 months prior to first dose). 6) presence of non-drug-induced psychosis through their life time. 7) unable to use a computer to the level requires to perform cognitive tasks.

### Design

This study was incorporated into a larger clinical trial examining the safety and tolerability of microdoses of LSD in healthy older adults. A randomised, double-blind, placebo-controlled design was used. Participants were randomly assigned to one of the four arms (12 per arm) that received either placebo or one of the three LSD doses (5μg, 10 μg or 20 μg). For drug manufacture information please see the previous study (Family et al., 2020).

#### Ignore /Update task

The task was a modified delay match-to-sample task designed to explore the effect of ignoring and updating on working memory (Fallon & Cools, 2014). The standard delay match-to-sample format was modified so that novel items were presented in the delay period between encoding items and probe (depending upon the condition; Figure 1). After encoding two memoranda (2000ms), there was a delay (4000ms) and participants were presented with either new memoranda (ignore and update) or a blank screen with a fixation cross (maintain). In the ignore condition, participants were presented novel stimuli that had to be ignored and prevented from entering working memory whereas in the update condition the novel stimuli had to be updated into working memory (displacing the previous memoranda as the target stimuli). Note, that participants were never explicitly instructed to ignore or update stimuli. but this was implicitly indicated by the presence or absence of the letter ‘T’, i.e., participants were told that they only had to remember items presented with the letter ‘T’. After a second delay period, participants were presented with a probe item and had to say whether the probe item matched or did not match the relevant target items (‘the last images presented with the letter ‘T’). Stimuli were randomly generated ‘spirographs’ (Fallon, Dolfen, et al., 2019)

**Figure 1.**
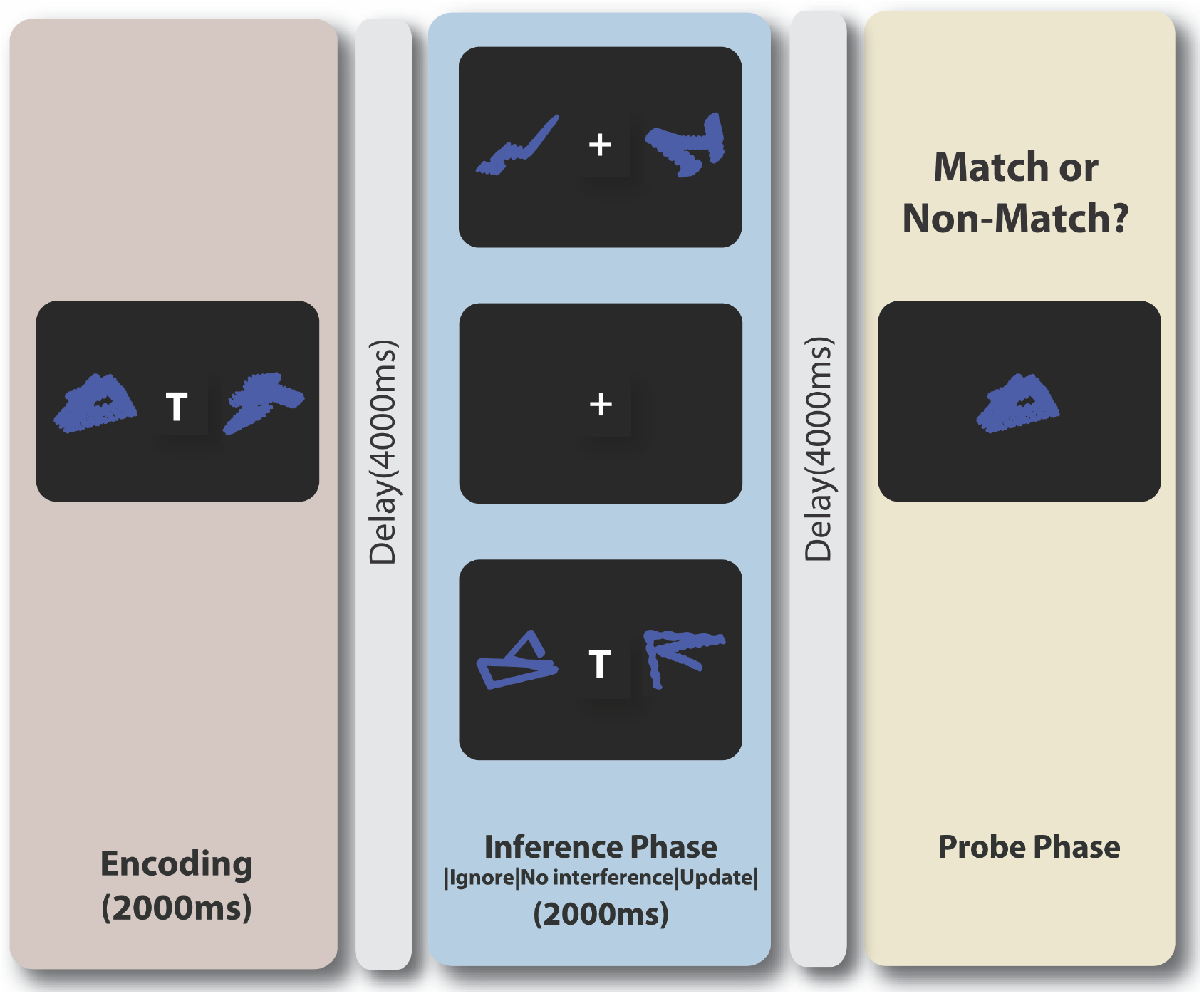
Ignore/Update task. The task was modified delay match-to-sample task. In these tasks, working memory is assessed at the end of each trial by asking participants to state whether a probe image matches or does not match the target stimuli. At the beginning of each trial. participants were required to encode two memoranda as the initial target stimuli. After a delay, and depending on the trial condition, they were then presented with either a blank screen or two novel memoranda. When presented with novel memoranda, these images had to either be updated into working memory as the new target stimuli (if a ‘T’ was present) or ignored completely (if no was ‘T’ present). Maintain trials featured a blank screen with no memoranda. Finally, participants were presented with a probe image and had to state whether this probe image matched or did not match the respective relevant target for each condition.

Participants completed 72 trials in total (24 trials for each trial type). Half of these trials were match trials and half of them were non-match trials. Non-match trials were equally drawn from novel memoranda and memoranda drawn from presented- but now non-target – stimuli.

### Analysis

The results were analyses using mixed general/generalized linear models using the fitlme/fitglme commands in Matlab (Matlab 2018). Accuracy (correct or incorrect) for each trial were examined across conditions and drug conditions in a generalized linear mixed model using the binomial distribution with logit link. For response latencies (correct trials only), a generalised linear model with the ‘normal’ distribution and ‘identity’ link were used. In both models, condition (ignore, maintain and update) and drug condition were incorporated as fixed effect dummy variables (‘effects’ option). Participant (split according to drug condition) were also entered as random effects (intercept).

## Results

### No evidence that LSD affects working memory accuracy or response latency

A generalised mixed model comparing accuracy rates for each condition (ignore, maintain and updating, descriptive statistics Table 1) across each drug condition found, as expected, a significant main effect of condition (*F*(2,3404) = 9.51, *p* <.0001). Participants were significantly less accurate for ignore trials compared to update trials (*F*(1,2273) = 12.97, *p* = .0003) and for ignore compared to maintain trials (F(1,2267) = 14.64, p = .0001). There was no evidence for a difference between update and maintain trials (F<1).

**Table 1.**
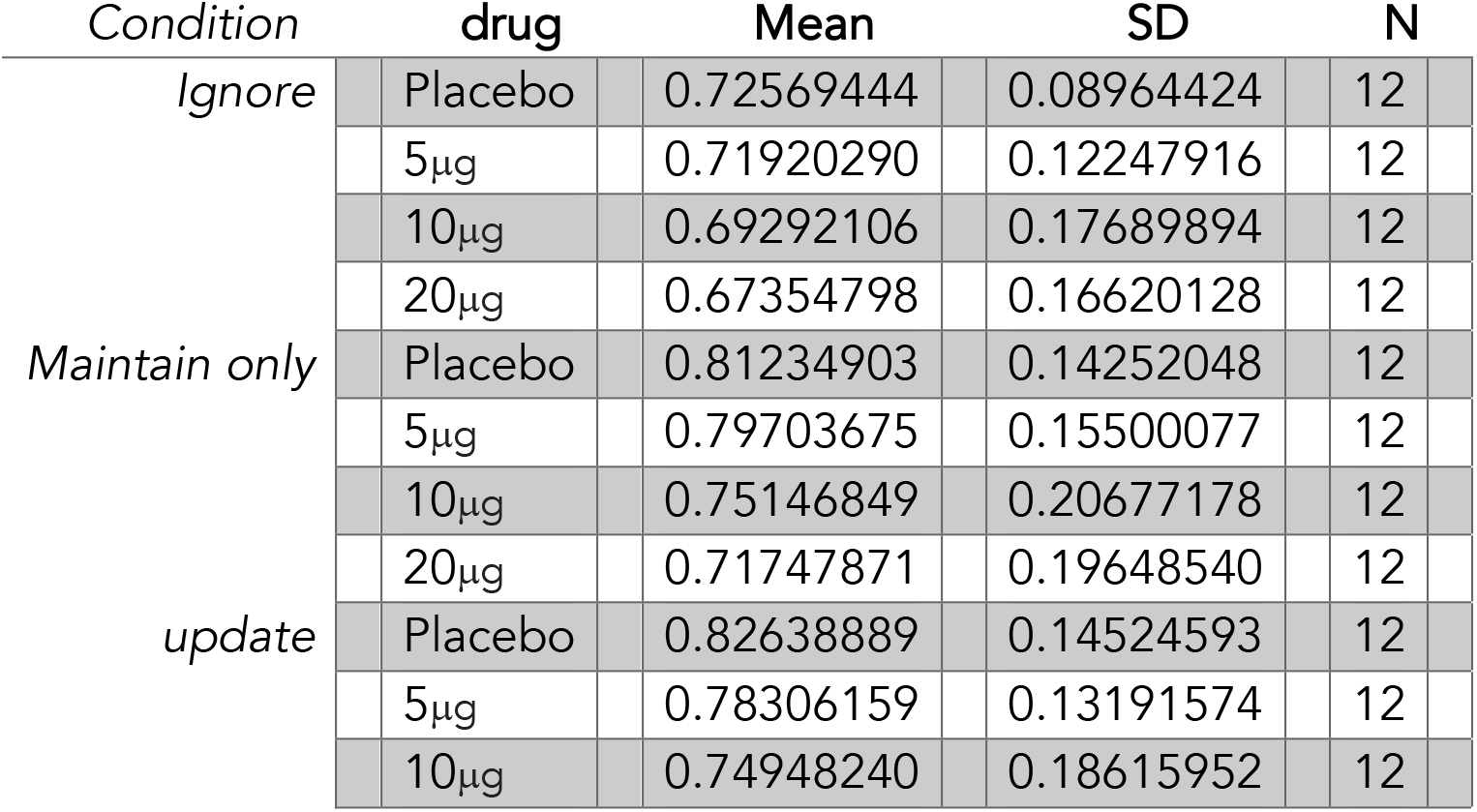
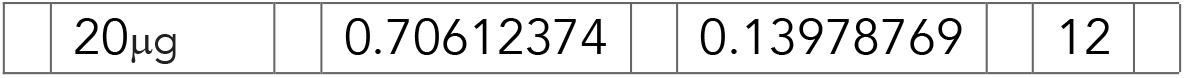
Means and standard deviations (SD) for accuracy for each task condition in each drug group.

There was no evidence for a significant effect of drug dose (*F*(3,3404) = 1.11, *p* = .34) or interaction between condition and drug dose (*F*<1).

**Figure 2:**
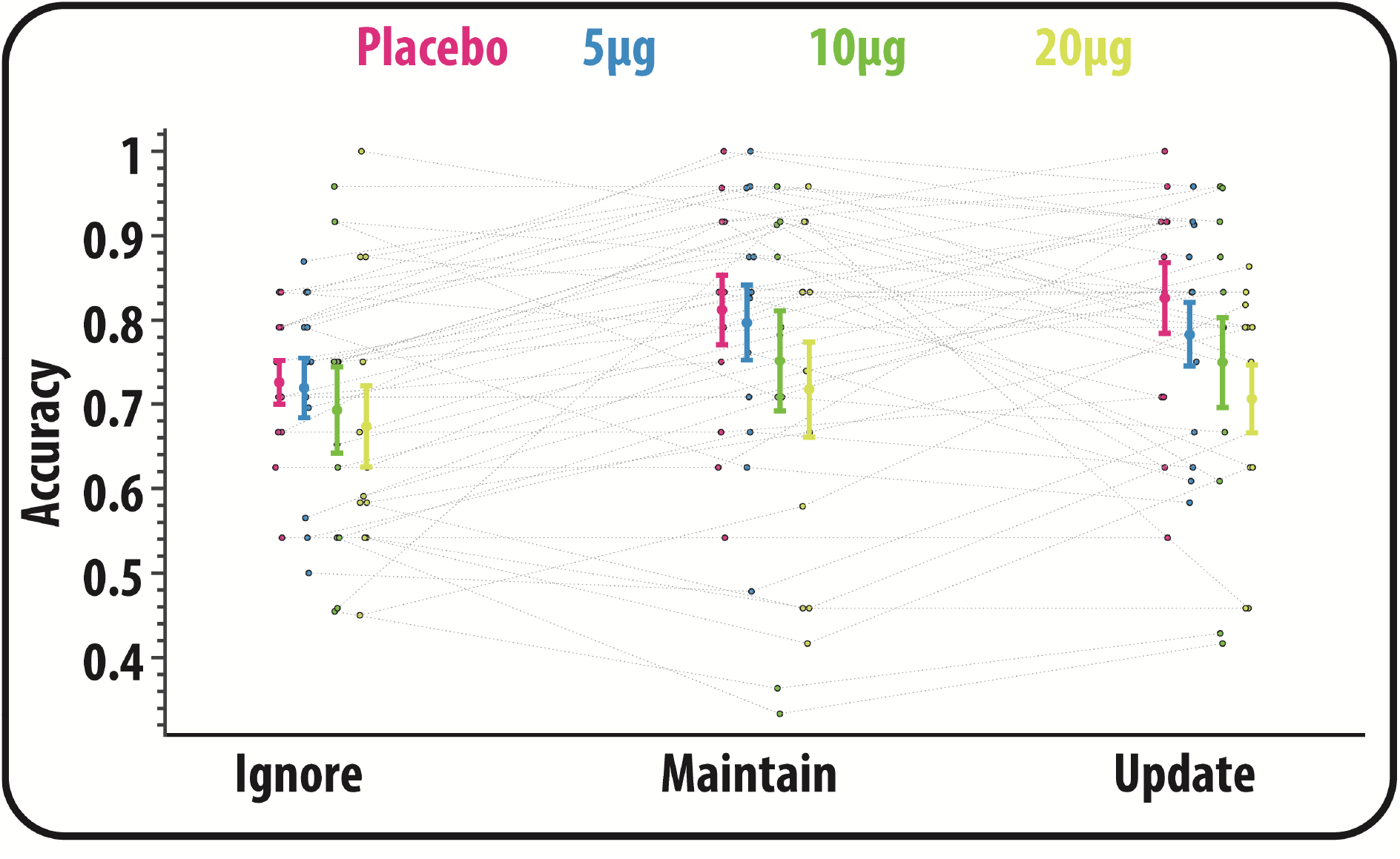
Accuracy rates according to condition and drug dose. Error bars reflect the standard error of the mean.

Similar results were obtained for reaction times (correct answers only). A significant main effect of condition was observed (*F*(2,2748) = 14.91, *p* <.0001). There was no evidence for an effect of drug dose or interaction between drug dose and condition (*F*s<1).

**Figure 2:**
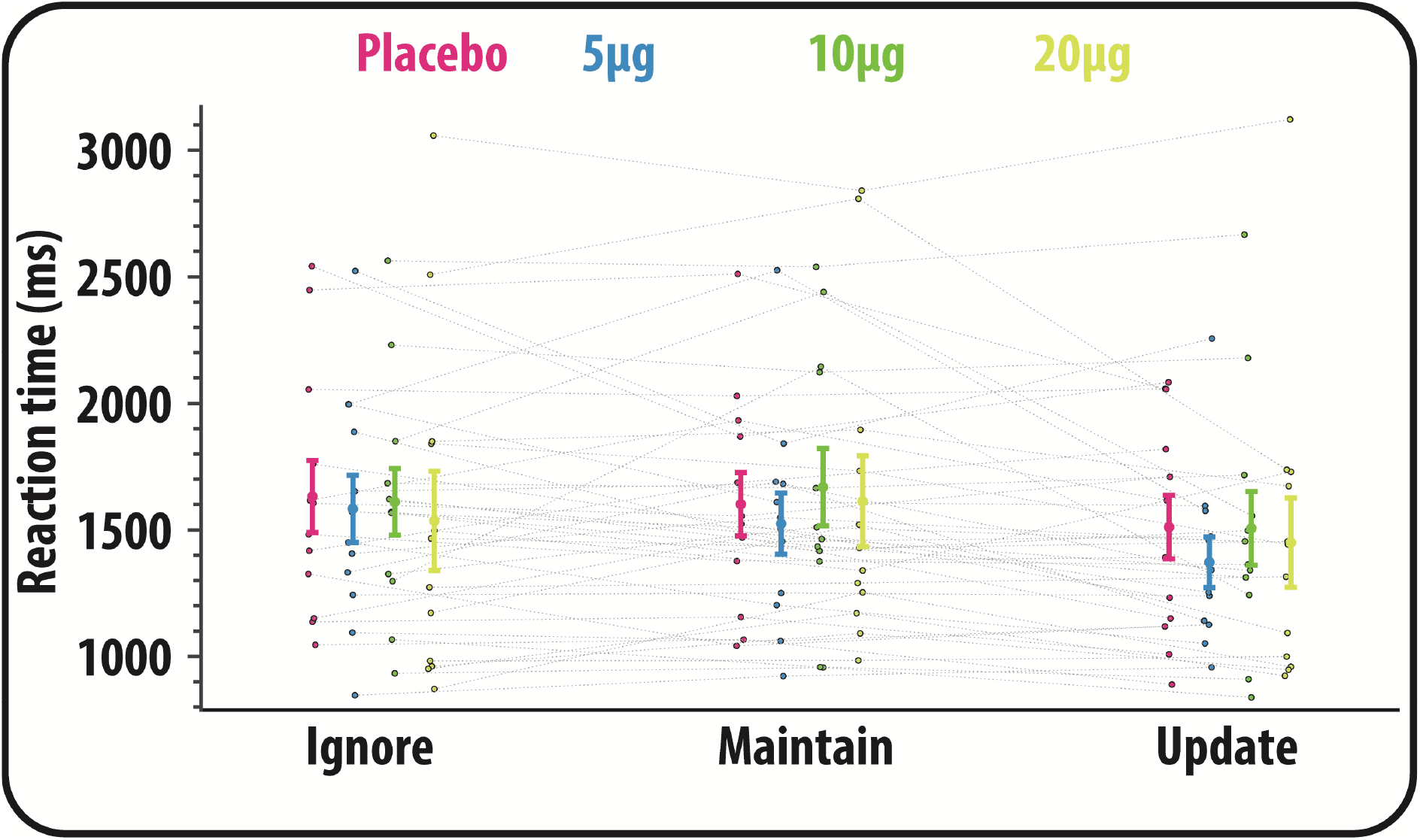
Reaction times (correct responses only) according to condition and drug dose. Error bars reflect the standard error of the mean.

## Discussion

Microdoses of LSD are frequently taken by people who wish to improve their cognitive or emotional wellbeing (Rifkin et al., 2020). This study examined whether very low doses of LSD affect overall working memory recall and whether there is any evidence that LSD exerts differentially effects on ignoring and updating information. Overall, there was no evidence to support the contention that LSD microdoses affect recall or the stability and flexibility of mental representations. Although these results should be regarded as preliminary given the small sample sizes, the lack of an effect of LSD on working memory in this study supports the results of previous studies that have failed to find mnemonic effects of LSD (Bershad et al., 2019).

LSD microdoses are thought to exert their psychological effects through acting on serotonin 5-HT_2A_ receptors or dopamine D_2_ receptors (De Gregorio et al., 2016). Given empirical and computational work (Cools & D’Esposito, 2011; Durstewitz & Seamans, 2008; Hazy et al., 2007), modulating signalling along either of these pathways could have been anticipated to affect working memory or the stability of mental representations. In this study, there was no evidence for this. Several factors could be responsible for this.

It could be argued that due to its small sample sizes the study was insufficiently powered to detect a significant effect of LSD microdoses on working memory. However, while the small sample size precludes any strong inferences from the lack of significant effects, other investigations from this study were able to detect effects of LSD microdoses on time perception (Yanakieva et al., 2019). Thus, it is possible that the effect of LSD microdoses on working memory is smaller than time perception (Yanakieva et al., 2019). Consistent with this, LSD microdoses in small samples have been found to exert bigger effects on emotion-related processing than on the n-back working memory paradigm (Bershad et al., 2019).

One objection to results of de Wit study that investigated working memory after LSD microdose is that the n-back task was insufficiently sensitive to detect an effect, i.e., that given that the n-back task cannot distinguish between the stability and flexibility of working memory, antagonistic effects of LSD on working memory sub-processes could have been missed. It is unlikely that the task used here was insufficiently sensitive to detect these changes as similar designs have been used to uncover the differential effects of methylphenidate on ignoring and updating (Fallon et al., 2017). Therefore, the effect of LSD microdoses on stability and flexibility may be negligible. However, it has been well established that the effects of pharmacological compounds on working memory can be baseline dependent (Cools & D’Esposito, 2011; Frank & O’Reilly, 2006), with traits such as impulsivity (Cools et al., 2007), genetic background (Naef et al., 2017) or working memory performance (van der Schaaf et al., 2013). Indeed, factoring individual differences in baseline working memory has been found to be essential for probing the differential effects of dopaminergic drugs on the stability and flexibility of mental representations (Broadway et al., 2018; Fallon, Kienast, et al., 2019; Jongkees, 2020) – though see (Fallon, Zokaei, Norbury, et al., 2016). Therefore, a fuller and more authoratative answer on the effects of LSD microdoses on working memory await further, larger studies that are capable of capturing individual differences in drug responses.

## Acknowledgements

Data preparation and collection for this study was funded by Eleusis Therapeutics Ltd. SJF received no financial remuneration or benefits from this study. SJF is funded by the NIHR Biomedical Research Centre at University Hospitals Bristol NHS Foundation Trust and the University of Bristol. The views expressed in this publication are those of the author(s) and not necessarily those of the NHS, the National Institute for Health Research or the Department of Health and Social Care.

